# PRRSV-2 variant classification: a dynamic nomenclature for enhanced monitoring and surveillance

**DOI:** 10.1101/2024.08.20.608841

**Authors:** Kimberly VanderWaal, Nakarin Pamornchainavakul, Mariana Kikuti, Jianqiang Zhang, Michael Zeller, Giovani Trevisan, Stephanie Rossow, Mark Schwartz, Daniel C.L. Linhares, Derald J. Holtkamp, João Paulo Herrera da Silva, Cesar A. Corzo, Julia P. Baker, Tavis K. Anderson, Dennis N. Makau, Igor A.D. Paploski

## Abstract

Existing genetic classification systems for porcine reproductive and respiratory syndrome virus 2 (PRRSV-2), such as restriction fragment length polymorphisms (RFLPs) and sub-lineages, are unreliable indicators of genetic relatedness or lack sufficient resolution for epidemiological monitoring routinely conducted by veterinarians. Here, we outline a fine-scale classification system for PRRSV-2 genetic variants in the U.S. Based on >25,000 U.S. open-reading-frame 5 (ORF5) sequences, sub-lineages were divided into genetic variants using a clustering algorithm. Through classifying new sequences every three months and systematically identifying new variants across eight years, we demonstrated that prospective implementation of the variant classification system produced robust, reproducible results across time and can dynamically accommodate new genetic diversity arising from virus evolution. From 2015 and 2023, 118 variants were identified, with ∼48 active variants per year, of which 26 were common (detected >50 times). Mean within-variant genetic distance was 2.4% (max: 4.8%). The mean distance to the closest related variant was 4.9%. A routinely updated webtool (https://stemma.shinyapps.io/PRRSLoom-variants/) was developed and is publicly available for end-users to assign newly generated sequences to a variant ID. This classification system relies on U.S. sequences from 2015 onwards; further efforts are required to extend this system to older or international sequences. Finally, we demonstrate how variant classification can better discriminate between previous and new strains on a farm, determine possible sources of new introductions into a farm/system, and track emerging variants regionally. Adoption of this classification system will enhance PRRSV-2 epidemiological monitoring, research, and communication, and improve industry responses to emerging genetic variants.

**Importance:** The development and implementation of a fine-scale classification system for PRRSV-2 genetic variants represents a significant advancement for monitoring PRRSV-2 occurrence in the swine industry. Based on systematically-applied criteria for variant identification using national-scale sequence data, this system addresses the shortcomings of existing classification methods by offering higher resolution and adaptability to capture emerging variants. This system provides a stable and reproducible method for classifying PRRSV-2 variants, facilitated by a freely available and regularly updated webtool for use by veterinarians and diagnostic labs. Although currently based on U.S. PRRSV-2 ORF5 sequences, this system can be expanded to include sequences from other countries, paving the way for a standardized global classification system. By enabling accurate and improved discrimination of PRRSV-2 genetic variants, this classification system significantly enhances the ability to monitor, research, and respond to PRRSV-2 outbreaks, ultimately supporting better management and control strategies in the swine industry.

## Introduction

In the U.S., Porcine reproductive and respiratory syndrome virus-type 2 (PRRSV-2) circulates within 30-50% of swine breeding farms in any given year (1, 2), causing both reproductive and respiratory impacts that result in >$600 million USD of productivity losses annually (3). These economic losses make PRRSV the most important virus affecting swine in the U.S. Classified as the species *Betaarterivirus americense* (the former species *Betaaterivirus suid 2*) in the family *Arteriviridae* and order *Nidovirales*, PRRSV-2 is a rapidly evolving RNA virus characterized by enormous genetic and antigenic variability in the U.S. and globally (4-6). Control of this virus is hindered by routine emergence of novel, sometimes more virulent genetic variants (7-9), which result in recurrent epidemic waves of viral spread in the industry (5, 10).

PRRSV-2 is also one of the most sequenced viruses in the world (11), largely because sequencing is used by animal health professionals as a tool for routine monitoring of virus circulation within and between farms. While phylogenetic analysis is still the gold standard for interpretation of sequence data, practitioners and field epidemiologists often find it faster and more convenient to have a name in which they can refer to a given genetic variant as part of everyday communication and outbreak investigations. Currently, the naming method used by the industry to discriminate between sequences is restriction fragment length polymorphism (RFLP)-typing (12), sometimes in combination with an additional label corresponding to phylogenetic lineage (4, 5). However, lineages and sub-lineages are large and diverse, and hence are too coarse for on-farm disease monitoring, and using RFLP-types to refer to PRRSV-2 viruses often leads to misleading conclusions (e.g., viruses assigned to the same RFLP-type often are not genetically similar, and vice versa)(13-15). For example, RFLP 1-4-4, which is one of the most abundantly reported in the U.S. today, occurs in seven different lineages (16).

In previous work, VanderWaal *et al*. (15) evaluated and compared 140 approaches for fine-scale classification of ORF5 sequences. Three methods were found to be robust and reproducible, and thus could form the foundation for fine-scale classification of PRRSV-2 below the sub-lineage level. However, previous work did not explore the performance of PRRSV-2 variant classification on a rolling basis, and it is necessary to validate the performance and associated procedures for fine-scale classification that accommodates expanding genetic diversity on a prospective basis.

Taking insights and needs of practitioners and diagnosticians alongside a rigorous comparison of alternative approaches for classifying PRRSV-2 (15), the purpose of this paper is to introduce a new fine-scale genetic classification system for PRRSV-2 that is tailored to meet the needs of animal health professionals. Specifically, we outline criteria used for defining PRRSV-2 genetic variants, establish and test procedures for prospective implementation of the system, and assess the adaptability of the classification system to accommodate expanding genetic diversity at national scales. We also introduce a machine-learning webtool that can be used to identify the variant to which newly generated sequences belong and introduce naming conventions for PRRSV-2 variants. Finally, we report the results of a survey conducted with field practitioners on their motivations for submitting samples for sequencing and demonstrate how variant classification can enhance the utility of sequence data for the purposes of epidemiological monitoring and surveillance.

## Results

### Variant classification

Utilizing sequences from the U.S. from 2015-2023, 25,403 PRRSV-2 ORF5 sequences were analyzed on a rolling quarterly basis to simulate prospective application of the variant classification system; each quarter, groups of closely related sequences were identified in phylogenetic trees using a clustering algorithm and defined as a variant if the group a) had five or more sequences, b) showed robust support of their shared ancestry in the ORF5 phylogeny (bootstrap value >85), and c) was >2% different than the nearest named variant. Any sequences belonging to clades that did not meet these requirements were labeled as “unclassified.” In total, the fine-scale classification system identified 118 genetic variants, 37 of which were common (detected >50 times) and 19 were rare (detected <10 times). 89.7% of sequences belonged to common variants, while 1.3% of sequences belonged to rare variants. The median number of sequences per variant was 25.5, with an interquartile range (IQR) of 11.25 – 68.75 sequences. The average within-variant genetic distance was 2.4% (IQR: 1.6 −3.2%, max: 4.8%). The mean distance to the closest related variant was 4.9% (IQR: 2.5 – 5.6%). Median bootstrap support for variants was 100 (IQR: 91.7 – 100). The distribution of variants on a phylogenetic tree is shown in Figure 1. Variant nomenclature incorporated the sub-lineage to which the variant belonged, followed by an integer (i.e., 1A.3 and 1H.3 are the third variants identified within sub-lineage 1A and 1H, respectively). For contemporary sequences (2015 onwards), several variants were a one-to-one correspondence with vaccine-like sequences, namely variant 5A.1 (Ingelvac PRRS MLV - Boehringer Ingelheim Animal Health, Duluth, GA), 8A.1 (Ingelvac PRRS ATP - Boehringer Ingelheim Animal Health, Duluth, GA), 8C.1 (Fostera PRRS - Zoetis, Parsipanny, NH), 1D.2 (Prevacent PRRS, Elanco, Greenfield, Indiana), 7.1 (PrimePac PRRS, Merck, Rahway, NJ), and 1F.1 (PRRSGard, Pharmgate Animal Health, Wilmington, NC).

**Figure 1.**
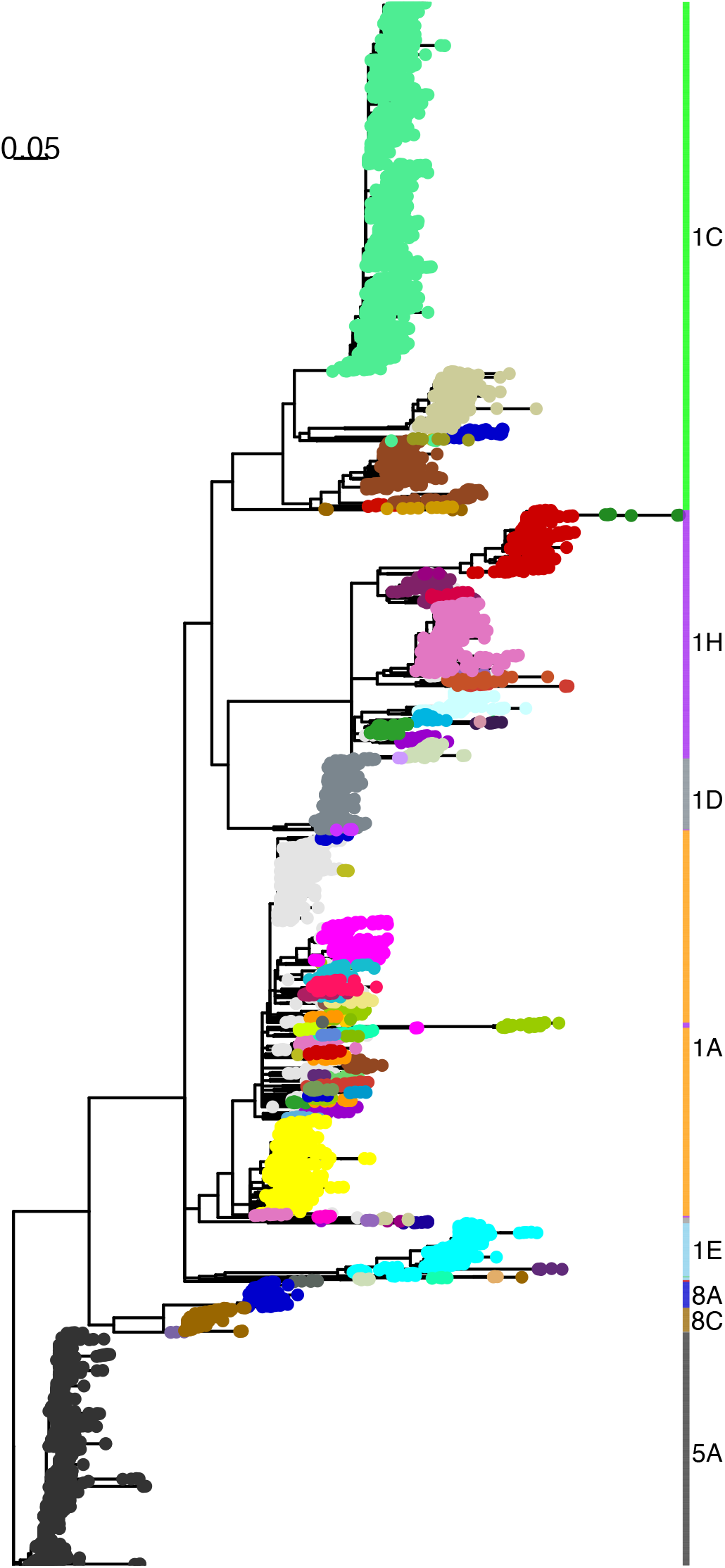
36-month phylogenetic tree at last timepoint (December, 2023). Tip colors indicate variant. Color bars indicate sub-lineage.

Across 21 quarterly datasets (each containing 36-months of data, 2015-2023), genetic diversity within a variant did not show an increasing trend through time (Supplementary Figure 1a). Clade purity was calculated for each variant as the proportion of sequences in a phylogenetic clade that were assigned to the same variant ID. Clade purity was consistently high across quarters, with a median of 100% (IQR: 99% – 100%, mean: 89.9%, Supplementary Figure 1b), indicating that variants formed compact groups and were not inter-mixed across the phylogenetic tree. Initially, ∼36% of sequences could not be reliably grouped into a well-supported variant clades and were considered “unclassified,” but this value reduced and stabilized to ∼11% in 2020 and 2021, and ∼7% in 2022 and 2023. Lineage 1A accounted for 94.6% of unclassified sequences. Lineage 1A has a lower genetic diversity than other sub-lineages due to its more recent emergence approximately 10 years ago (5), and classification for this sub-lineage improved as clades became more diverged through time.

The median number of active variants per year was 48 (Figure 3). This compares to 65 and 112, respectively RFLP-types and Lineage+RFLP, which are currently employed for fine-scale PRRSV classification. The median number of “common” active variants was 26, 25, and 39 for variants, RFLP-types, and Lineage+RFLPs, respectively. Thus, the new classification system does not result in a greater number of IDs than the industry currently is accustomed to with RFLP-types.

There was a median of 19 new variants per year, but only 4 new common variants (those that would eventually be detected >50 times), demonstrating that variant classification is able to scale-up to accommodate newly emerging PRRSV diversity (Figure 2). In contrast, there were no new common RFLP-types across the study period. Most newly-identified variants were created from sequences that were previously “unclassified”, and not from splits of existing variants. In total, 0.9% of sequences were re-named (i.e., a result of splitting a variant) at some point during the 21 quarters assessed here.

**Figure 2.**
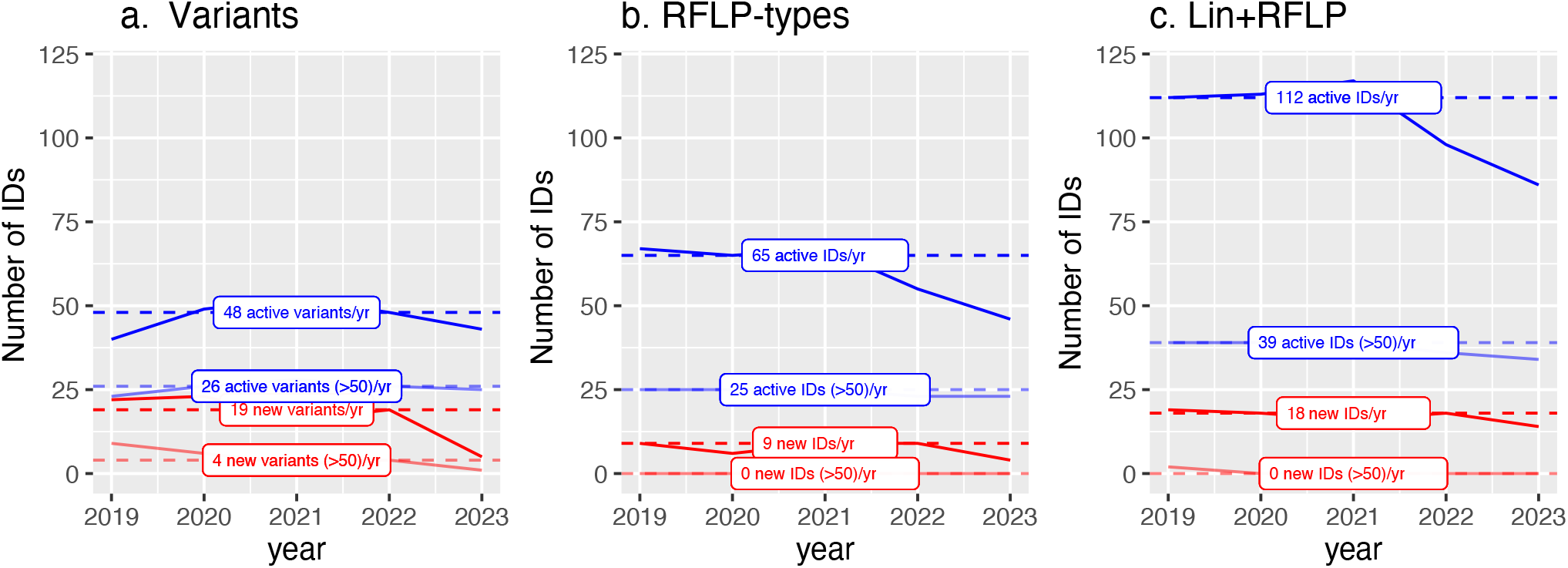
Number of variants per year. Yearly number of active (blue) and new (red) variants for each classification method, with those that reach at least 50 sequences considered “common.” Solid lines show the number per year. Dashed lines show the median number across years.

Using a subset of data, we also constructed time-scaled phylogenetic trees to contextualize variant emergence and divergence on a timeframe that is interpretable for epidemiological investigations of within- and between-farm transmission. We found that sequences belonging to the same variant typically descended from a common ancestor that existed ∼2.3 years prior (median: 2.3 years, IQR: 1.5 – 3.5 years), which can be interpreted as that all sequences belonging to a single variant were part of the same chains of transmission originating ∼2 to 3 years prior. This gives a timeframe for which to search for epidemiological connections amongst cases. Divergence time from the closest relatives was 3.9 years (median: 2.9 years; IQR: 3.1 – 4.6 years). Clade purity in time-scaled trees was high, with a median of 1.0 and interquartile range from 90 – 100%.

### Tools for assigning variant IDs to new sequences

Across the 21 quarters, the quarterly-updated assignment algorithm had accuracies ranging between 97.1 - 99.9%, groupwise accuracies between 95.1-99.8%, mean precision (positive predictive value) between 96.6-99.9%, and mean recall between 95.1-99.8%. The percentage of sequences that were undetermined (i.e., probability of assignment was <0.25) ranged from 0.2% - 8.1%, with a median of 4.4%. The most up-to-date model is accessible via a RShiny webtool (https://stemma.shinyapps.io/PRRSLoom-variants/) and the trained model is available on Github in both R and Python (https://github.com/kvanderwaal/prrsv2_classification). Whether using the webtool or the R/Python code, the user uploads ORF5 sequence/s in fasta format, which are then realigned to the PRRSV-2 prototype sequence VR2332 (Genbank accession EF536003). Sequences are not saved or retained by the webtool in any way. The tool then estimates the probability that the sequence belongs to each defined variant. For each sequence, outputs include the assignment probability for the variant ID with the highest (top) and second highest probability. A final assignment is also given, with sequences that could not be assigned to any variant with >0.25 probability listed as “undetermined.” It is also possible that the variant with the highest probability is not substantially greater than the second highest, which may indicate potential misclassification. If the highest probability is more than double the second highest probability, then the assigned variant ID can be interpreted with greater confidence. While this paper reports results up to December 2023, the PRRSLoom-variant shiny application and the pre-trained model has been updated to reflect recent variant classifications, and this will be maintained on a quarterly basis.

### Survey on use of PRRSV-2 sequence data by animal health professionals

In a survey administered by the American Association of Swine Veterinarians, swine practitioners (n = 92) were asked to rank the primary motivations for which they submitted samples for sequencing. The motivations that were consistently highly ranked included 1) *Anticipate and track the spread of novel and emerging variants*, 2) *Discriminate between previous and new wild-type strains on the same farm*, and 3) *Determine possible source of introduction* (Figure 3).

**Figure 3.**
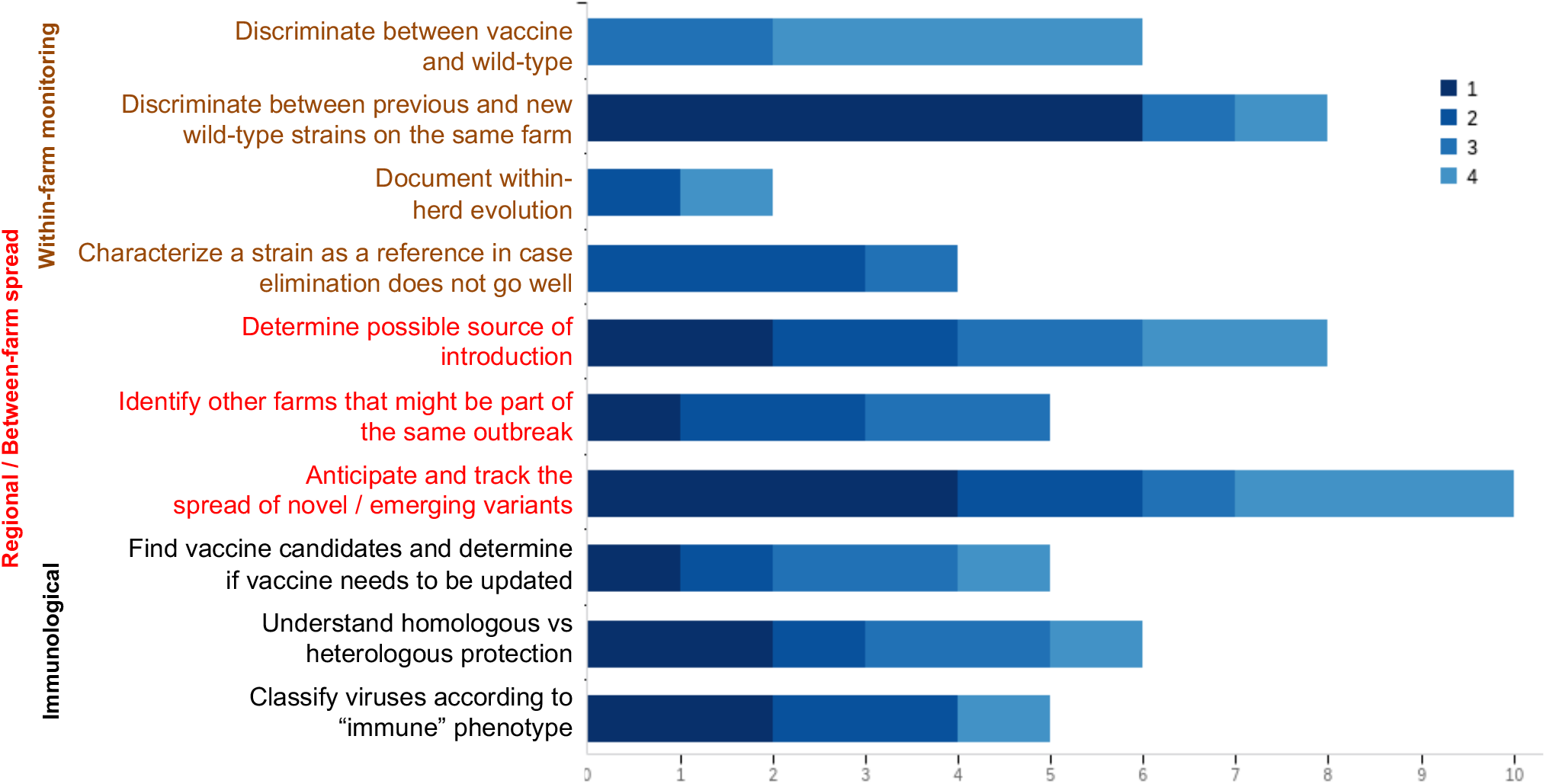
Results of survey where animal health professionals were asked to rank their top four reasons for submitting samples for sequencing from a list of 10 options. Bars represent the number of respondents that selected each answer, with color shading representing rank (with 1 being high).

1) *Anticipate and track the spread of novel and emerging variants:* To better visualize how the new classification system tracks the spread of emerging variants, we selected variants that had <25 sequences at the time of naming and >200 sequences by the end of the study period. Five variants met these criteria (Figure 4). We also considered the 1H.18 variant, due to interest in this variant at the time of writing (17). There was a median of 10.5 different RFLP-types per variant. Of the common RFLP-types (n>50), none were exclusive to any of the emerging variants. Indeed, these common RFLPs were all found in ≥3 of the six emerging variants, and across a median of 33 variants overall. These insights show the benefits of using variant classification as opposed to RFLP-typing for identifying and tracking emerging strains of the virus.

**Figure 4.**
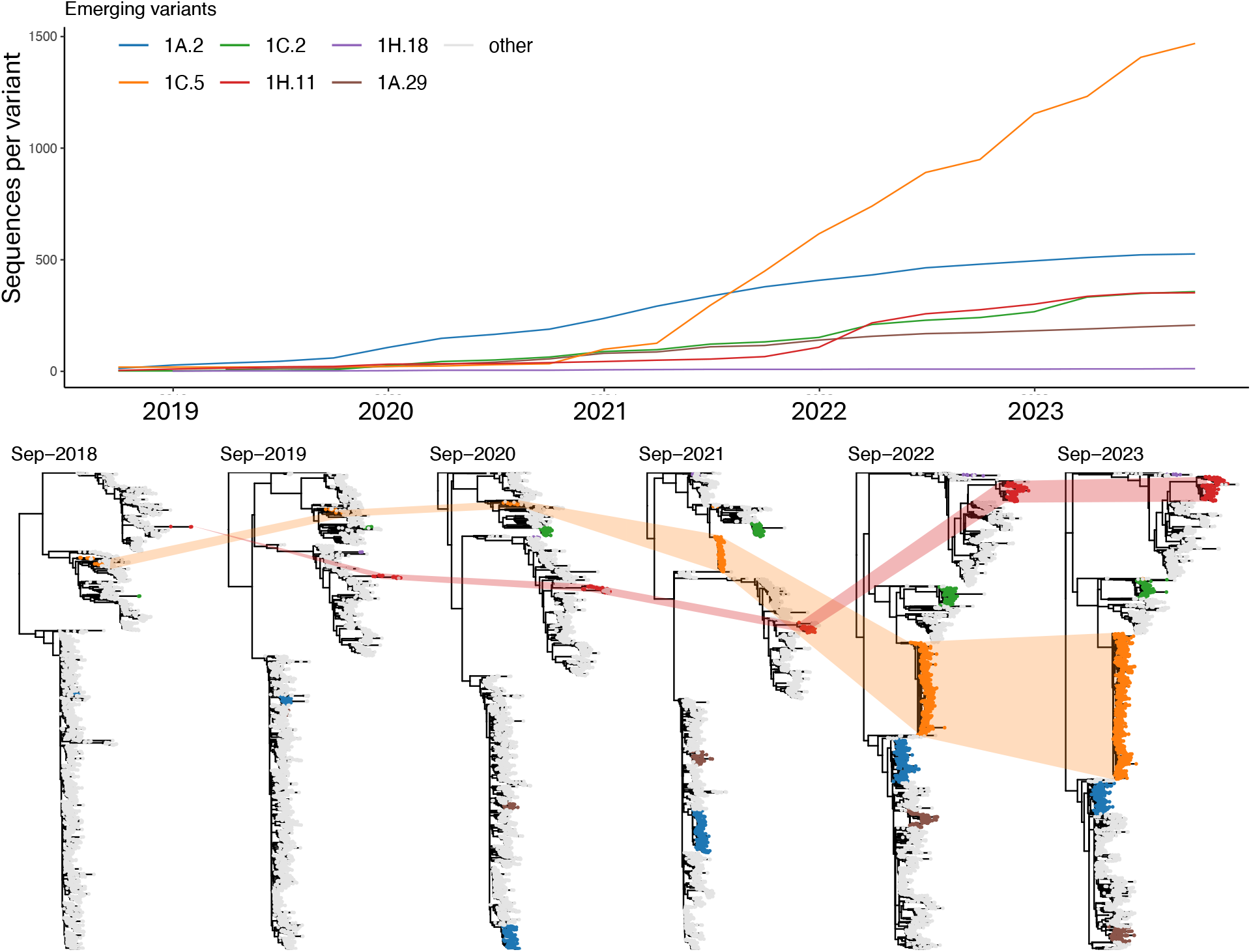
Cumulative number of sequences per emerging variant over time (top). Phylogenetic trees (bottom) from September of each year, with emerging variants colored and all other sequences shown in gray.

To 2) *Discriminate between previous and new wild-type strains on the same farm*, and 3) *Determine possible source of introduction*, we partnered with production system veterinarians and applied the new variant classifications to sequences collected from their farms. System 1 shared 28 PRRSV-2 ORF5 sequences from 12 farms, with a particular interest in 13 sequences from Farm 5, which was a sow farm (Figure 5). This farm experienced four PRRSV circulation events: variant 1B.8 in 2015-2016, 1C.3 in 2016-2018, 1A.13 in 2019, and 1C.5 in 2020-2024. Of note, RFLP-types or lineages were not able to discriminate between new introductions on Farm 5. Either a new introduction did not receive a unique label (as in 2020, where sub-lineages failed to discern a new 1C virus, despite having <91% nucleotide identity with the previous 1C virus on the farm), or multiple sequences that were part of these same circulation event received different labels (three different RFLP-types amongst the five 1C.3 sequences, despite having >98% nucleotide identity). This limitation of RFLP-types is more thoroughly quantified in VanderWaal *et al*. (15), wherein 43% of on-farm circulation event attributable to a single variant had multiple associated RFLP-types.

**Figure 5:**
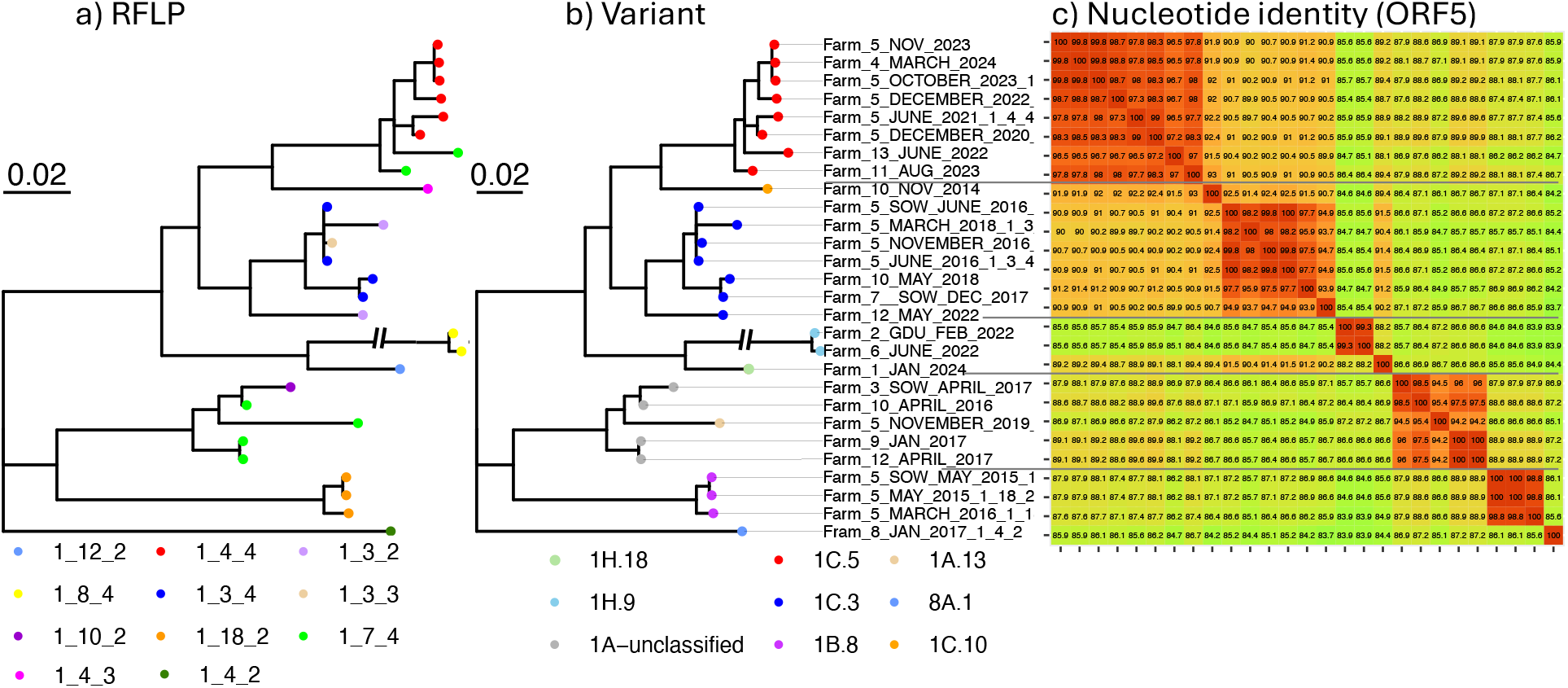
Phylogenetic tree of ORF5 sequences for System 1 reconstructed by Bayesian inference (mrBayes v3.2 (18)), rooted on the only sequence not belonging to lineage 1 (sub-lineage 8A), with tip color indicating a) RFLP-type, and b) variant. c) ORF5 nucleotide identity, with redder colors representing higher identity.

System 2, which is a large production system operating in five states, shared 1,095 ORF5 sequences from 2014 to 2022. In the phylogenetic tree in Figure 6, the majority of the sequences belonged to sub-lineages 1C, 1H, and 1E. Lineages and sub-lineage do not provide sufficient resolution to distinguish new and already circulating wild-type viruses on farms, and also fail to provide a distinguishing label for one clade that contains recombinant viruses. The vast majority of sequences belong to two RFLP-types (1-8-4 and 1-4-4), which do not cluster together genetically. Furthermore, the RFLP 1-4-4 sequences are not related to the recently emerged outbreak variant bearing the same RFLP-type (the so-called novel L1C-1-4-4 variant, which is referred to as 1C.5 in the new variant system (7)). In contrast, the variant classifications are aligned with clades that are visually well-differentiated and also provide a unique variant ID for the recombinant clade. Thus, the improved labeling of closely related sequences in the variant system enhances a practitioner’s ability to track spread between farms, detect the introduction of new variants into a farm or flow, and narrow the possible sources of introduction.

**Figure 6.**
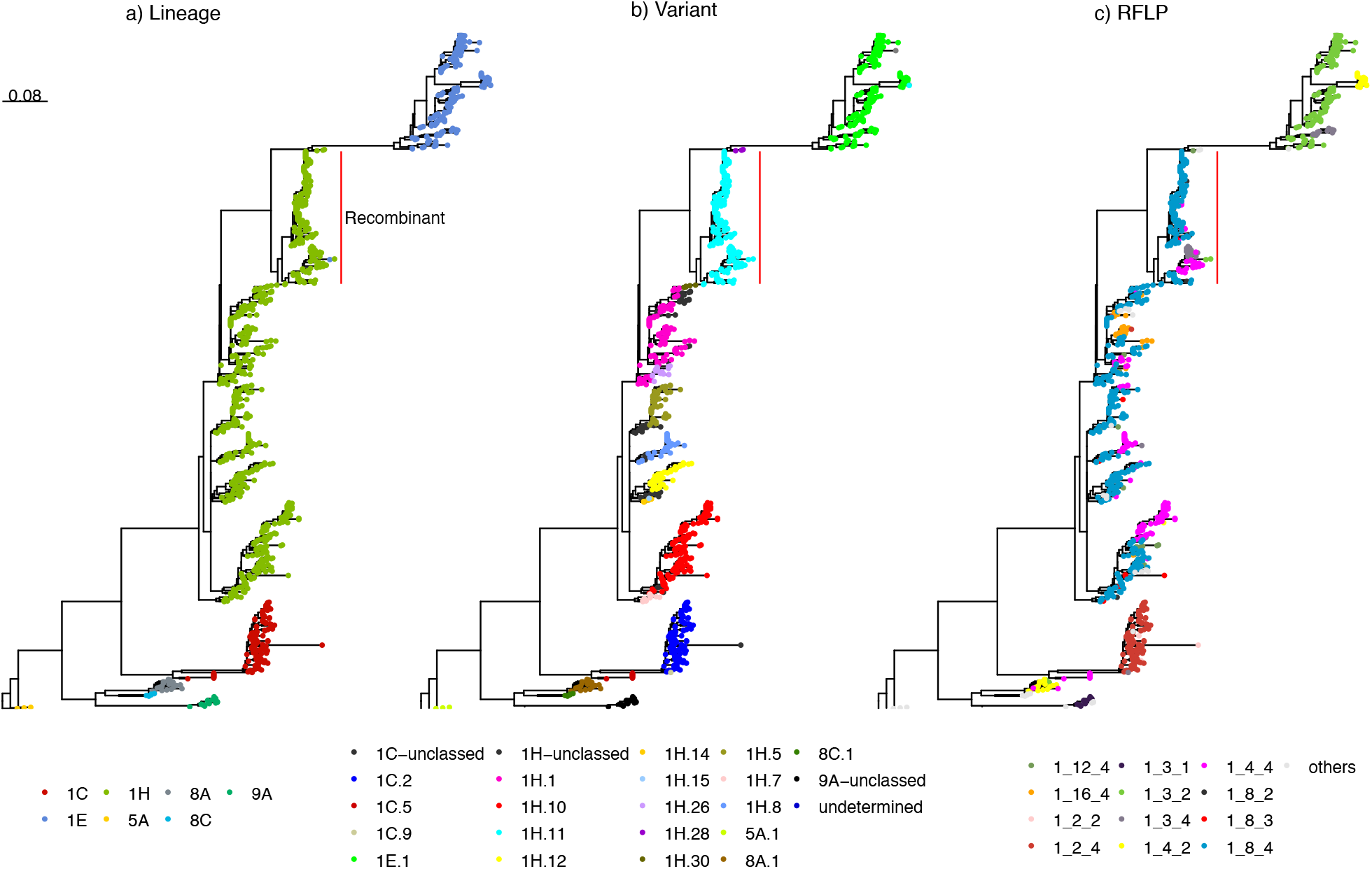
Maximum-likelihood tree of PRRSV-2 ORF5 sequences for System 2, colored by a) lineage, b) variant, and c) RFLP-type. The red bar indicates a clade that includes numerous recombinant sequences.

## Discussion

While the shortcomings of existing PRRSV-2 ORF5-based classification systems have been apparent as early as 2011 (19), recent advances in computational power and the creation of national-scale sequence databases have created the opportunity to finally address these issues. Here, we outline a fine-scale classification system (below sub-lineage level) for PRRSV-2 in the U.S. that is expandable to new genetic diversity that emerges as consequence of virus evolution. We lay out procedures for quarterly updating of the classification system and for assignment of newly generated sequences via a centrally-maintained machine learning model, which facilitates a unified naming scheme across the U.S. The level of granularity represented by genetic variants was tailored to meet the needs of animal health professionals, who primarily reported using sequence data for epidemiological monitoring. In this paper and in VanderWaal *et al*. 2024 (15), we demonstrate that as compared to RFLP-typing, variant classifications more reliably group viruses based on relatedness in the ORF5 gene, and provide better discrimination between unrelated viruses. This facilitates on-farm monitoring, detection of new introductions to a farm or production system, and tracking of regional between-farm spread.

Our fine-scale classification system is an extension of the lineage and sub-lineage classification first proposed in 2010 (20) and refined in the past five years (4, 5, 10). Lineages represent the broadest classification, with genetic distances typically <11% within a lineage based on ORF5. Sub-lineages typically have genetic distances <8.5% within the sub-lineage, and are made up of numerous genetic variants. Sequences belonging the same variant typically have an average genetic distance of 2 to 3% but can sometimes be as much as almost 5% different. Our intent is not to replace lineages, as we do believe that these larger classifications are useful for explorations of phenotype as well as tracking the macro-evolutionary dynamics of PRRSV-2. Therefore, we incorporate lineage into the IDs utilized in the variant classification system to provide a general zip-code of the variant within the larger genetic diversity of PRRSV-2.

A major advantage of a unified variant classification system within the U.S. is to facilitate communication amongst animal health professionals, diagnostic laboratories, and researchers. In the past, tracking of emerging variants was typically accomplished either by using RFLP-types or by calculating the genetic distance to “anchor” sequences established for a particular strain. RFLP-typing, with its associated limitations and inaccuracies, can generate confusion in the field about which farms are part of an outbreak, both missing farms that should be included (i.e., a closely related virus with a different RFLP-type) as well as sparking false-alarms (i.e., a distantly related virus with the same RFLP-type), as happened in the early days of the emergence of the 1C.5 variant (7). Calculating the genetic distance between sequences is a viable alternative for determining relatedness, but requires someone to set anchor sequences for a particular outbreak (which is usually only done for variants of heightened concern), and importantly, requires a several step process of sharing sequences (which are often considered confidential), aligning them and calculating distances in bioinformatic software. These steps are not required if the variant ID of the respective sequences is already assigned by diagnostic labs as part of their reports to clients.

However, an important caveat is that variant classification is not based on immunological or virulence variability (i.e., the phenotype) of the virus, and most variants will not have been fully characterized from a phenotypic standpoint even when a whole genome is available. Thus, variant classification is not designed to provide information on the clinical manifestations of the virus in a herd, which is influenced by a myriad of factors external to the virus itself (e.g., co-infections, host genetics, immunological history, etc.)(21-24). Variant classification also does not directly translate to immunological cross-protection. Although viruses labeled as the same variant are more genetically homologous on ORF5, cross-protection is not simply a function of genetic distance between viruses (25, 26). Furthermore, whole genome data is required for phenotypic investigations. That being said, variant classification may be too fine-scale to expect major phenotypic differences amongst closely related variants. However, if we made variants less granular, we would lose their utility for epidemiological investigations, such as determining possible sources of introduction and tracking regional spread.

Variant classification also facilitates the generation, organization and findability of additional information or research related to a particular variant, such as regional incidence trends, whether the group includes recombinant viruses, production impacts, etc. This, combined with the ease of cross-communication across diagnostic labs, researchers, and the field, could lay the foundation for additional research on PRRSV epidemiology, virology, and immunology.

Limitations to the proposed variant classification include the coverage of our sequence dataset. This system was based on U.S. PRRSV-2 sequences from 2015-present. Given that our dataset covers >55% of U.S. swine production, we believe that our data is reasonably representative of PRRSV-2 diversity circulating in the major pork producing regions in the country since 2015. Thus, we urge potential users of the webtool to be cognizant of the year of sequence collection and origin (country) of any sequences they may upload. While earlier sequences can be input into the webtool, they are likely to be predicted as “unclassified” given that diversity present in previous decades was not represented in our dataset. Similarly, it is not meant to encompass genetic diversity from other countries. Our analysis of time-scaled trees suggests that sequences belonging to the same variant typically evolved from a common ancestor that existed 1.5 to 3.5 years prior. This short timescale and the rapid evolutionary rate of the virus (5) supports the idea that variants circulating in the U.S. are likely distinct from variants in other countries, except perhaps in cases where there is very frequent transboundary movement (such as Canada). The system could be expanded to PRRSV-2 in other countries by incorporating their data into our quarterly updates to identify and name new variants. In the absence of sharing of larger databases, five representative sequences from a particular clade (either in the U.S. or elsewhere) that is currently unclassified and for which a variant ID is desired can be submitted to our system. Alternatively to using our platform, we suggest that other countries could adopt our criteria for defining a variant so that this term can be used more consistently across continents.

While having an improved naming scheme for PRRSV-2 genetic variants will not solve PRRS in the U.S., a classification system for field-based epidemiological monitoring is needed and has been requested by practitioners for many years. In this paper, we outlined the definition of genetic variants, the procedures for systemic identification of variants in a nationwide sequence dataset, and a validated workflow for routinely updating variant classification on a quarterly basis. The latter will ensure that the nomenclature system can dynamically adapt to evolving PRRSV-2 diversity across time and space. Variant classification will facilitate communication about outbreaks, tracking of emerging and endemic variants across time and space, as well as provide a framework to more rigorously analyze the genetic basis of variability in phenotype or production impacts. Finally, this work was conducted with iterative feedback from a working group of veterinarians, researchers, and diagnosticians. This close engagement with stakeholders and end-users has been crucial for the operationalization and adoption of the variant classification, ensuring that it is tailored to the needs of animal health professionals utilizing sequence data for decisions on disease management in the field.

## Methods

### Data source and pre-processing

Sequence data were obtained from the Morrison Swine Health Monitoring Project (MSHMP), which is a voluntary initiative operated by University of Minnesota that monitors PRRS occurrence in the U.S. MSHMP was initiated in 2011, and currently collects sequence data for farms belonging to 37 production systems, accounting for >55% of the U.S. pig population (2). Participating production systems share PRRSV ORF5 sequences that are generated as part of routine monitoring and outbreak investigations in breeding, gilt developing units, growing and finishing herds (27). Sequences are generally obtained either directly from each MSHMP participant or from the main veterinary diagnostic laboratory where participants submit their diagnostic samples. Meta-data for each sequence include farm name, date and farm type of origin (e.g., breeding or growing herd).

16,260 sequences were available from October 1, 2015 to June 30, 2021. These sequences were used to establish the rolling procedures for updating the classification system across time. An additional 9,143 sequences were available from July 1, 2021 to December 31, 2023. Based on VanderWaal *et al*., sequence datasets that lack duplicated (100% nucleotide identity) sequences produced the most consistent variant classifications (15). Therefore, sequences with 100% identity were de-duplicated before phylogenetic tree building. Duplicated sequences were retained for calculations of the frequency and mean genetic distances of variants. Sequences with ≥4 ambiguous nucleotides (0.5% of ORF5) or with gaps greater than 24 positions were removed from the dataset (4). To assess how the system would function if utilized prospectively, we initiated the classification system in 2018 with 36 months of data (2015-18), then added new data every three months up until December 2023. Thus, each quarter of data included the previous 36 months of sequence data, with a median of 8,620 sequences per set.

### Tree building

For each 36-month dataset, sequences were aligned to a consensus reference sequence based on all previous data with --6merpair, --keeplength, and --addfragments options of the MAFFT algorithm (28, 29). All tree-building unless otherwise specified utilized the maximum likelihood method performed using IQ-TREE2 with 1,000 ultrafast bootstraps (30). As described in VanderWaal *et al*. (15), strict majority-rule consensus trees were constructed (clades with bootstrap support <50 were collapsed), with the general time reversible substitution model with empirical base frequencies and gamma plus invariant site heterogeneity (GTR+F+I+G4). The *ggtree* package in R was used for all tree visualizations, with trees re-rooted on Lineage 5, which contains the PRRSV-2 prototype virus (VR2332, GenBank accession number EF536003)(20, 31).

### Variant classification: Initialization

#### Initial timestep

Utilizing the first 36-month tree (9783 sequences October 1, 2015 – September 30, 2018), a tree-based clustering approach was applied to the phylogeny using the average-clade method in *TreeCluster* package available in Python (32); clusters of genetically related sequences in the trees were referred to as “variants.” Briefly, this method identifies monophyletic clades where the average pairwise patristic distance between sequences within the clade is <7%. This threshold was selected based on a rigorous comparison of thresholds performed by VanderWaal *et al*. (15). In that analysis, the observed average pairwise distance between sequences belonging to the same variant was 2.3% (15).

Additional steps were applied based on preliminary results showing that some variants defined on the first 36-month tree did not consistently group together in subsequent trees. First, post-processing of *TreeCluster* outputs was performed to merge clusters with low-support (Supplementary text). Second, additional criteria for defining a variant were that the group must have a) five of more sequences, b) robust support of their shared ancestry in the ORF5 phylogeny (bootstrap value >85 in the tree), and c) that the genetic distance to the nearest named variant must be >2%. Any sequences belonging to clades that did not meet these requirements were labeled as “unclassified.” Nomenclature for variants was the sub-lineage to which the variant belonged, followed by an integer (i.e., 1A.3 and 1H.3 are the third variants identified within sub-lineage 1A and 1H, respectively). To align with the five sub-groups within sub-lineage 1C delineated by (4), we identified the variants that corresponded to those groups and utilize the same IDs and continue numbering onward from 1C.6.

### Variant classification: Updating

With each new quarter, new sequences from the most recent 3 months (median: 663 sequences) are assigned to variants using the assignment algorithm trained at the end of the previous quarter (see next section). A new 36-month tree is constructed, and variant IDs annotated to the tree. The tree is systematically examined for new variants using *TreeCluster* as well as for splits in existing variants (see Supplementary text for details). Splits were systematically considered for variants where the 95^th^ percentile of pairwise genetic distances was ≥5% (based on sequences from the previous 12 months to better capture recent genetic divergence). A new variant was only created if a clade met the following conditions: 1) consisted of five or more sequences, 2) had robust support of their shared ancestry in the ORF5 phylogeny (bootstrap value >85 in the tree), and 3) that the genetic distance to the nearest named variant was >2%. If the creation of the new variant was due to a split in an existing variant, a new variant was only created if the minimum and median genetic distance between the new and original variant was >3 and >5%, respectively. These high thresholds were set to minimize the number of sequences being re-named as a result of variant splitting, as per the request of diagnostic laboratories. New variants receive names in the same manner as described above (i.e., if 1A.3 if split, one daughter group retains the name 1A.3, the other receives the next integer in the series, for example, 1A.8).

### Algorithm for assignment of new sequences

For prospective application of any classification system, it is desirable to be able to assign new sequences to variants without performing computationally heavy analysis. We thus trained a random forest machine learning algorithm to assign new sequences to the appropriate variant ID (15, 33). Up to 120 sequences per variant (approximately 10 per quarter) were randomly selected from the initial time step to build a training dataset, which was then appended quarterly with new sequence data. Using the training dataset for each quarter, a random forest algorithm was fitted using the *caret* package in R using ten-fold cross-validation and auto-tuning of the *mtry* hyper-parameter (34). In parallel, we also trained a random forest in Python for Python end-users (Supplementary text). Model performance on the training set was assessed using ten-fold cross-validation (i.e., performance evaluated on 10% of observations that were left out of 10 iterative random forest runs). We report the overall accuracy (percent of sequences correctly classified by the algorithm) for the training dataset. We also calculated the mean groupwise precision (*a*.*k*.*a*. positive predictive value), recall (a.k.a. sensitivity), and accuracy (i.e., percent of sequences correctly classified per variant was first calculated, and then a mean of these groupwise accuracies was reported).

Outputs from the trained algorithms include the probabilities of the first, second, and third most likely variant ID for a given sequence, with the highest probability ID being assigned to the sequence for downstream analyses of predictive performance. In some cases, the highest probability ID was quite low, indicating that the model had poor confidence in the assignment. Therefore, sequences with assignment probabilities of <0.25 were considered “undetermined,” and not considered in calculations of model accuracy. The proportion undetermined was tracked and reported. More stringent thresholds do not markedly improve model accuracy, but resulted in a higher percentage of undetermined sequences (15). The training dataset and algorithm are updated each quarter to include sequences (up to 120) from new variants as well as additional recent sequences from existing variants (up to 60). Older sequences and variants are not removed from the assignment algorithm, in order for the model to retain the ability to predict on older sequences from 2015-present. Only sequences with assignment probabilities of >0.4 were included in the training dataset.

An RShiny web-tool was developed and is updated quarterly (https://stemma.shinyapps.io/PRRSLoom-variants/) so that end-users can assign new sequences to variants. The updated algorithms are also available as R and Python scripts so that they can be used in command line or ported to external applications maintained by diagnostic laboratories or other groups (https://github.com/kvanderwaal/prrsv2_classification). This ensures that all potential end-users will obtain the same variant classifications, regardless of the platform.

### Genetic characterization and phylogenetic properties of variants

At the final timepoint, genetic characterization of variants produced by each approach included a) the number of variants identified, b) the number of “common” variants (n >50 sequences belonging to the variant), c) median size (sequences per variant) and interquartile range, d) percent of sequences belonging to common variants, e) percent of sequences belonging to rare variants (n <10 sequences), f) median bootstrap value and interquartile range of the ancestral node, and g) mean genetic distance (raw p-distance) within a variant. Finally, we calculated the h) genetic distance to the most closely related cluster for each variant.

Across all timepoints, we also evaluated the mean genetic distance through time to better understand how the mean within-variant distance may expand as a result of ongoing evolution, as well as clade purity over time to assess the tendency of sequences belonging to the same variant to remain grouped together in the tree over time. Clade purity was calculated as the proportion of sequences in a phylogenetic clade that were assigned to the same variant ID (See supplementary text for details).

We also calculated the number of new variants detected per year and number of active variants per year. The number of new variants per year was based on the calendar year of the earliest sequence belonging to a variant, and active variants per year included all variants whose earliest and latest detected sequences occurred before, during, or after the considered calendar year. For comparison purposes, these values were compared to RFLP-types and Lineage+RFLP types.

Using a subset of data, we also constructed time-scaled phylogenetic trees (Supplementary methods) to contextualize the timeline of variant emergence and divergence on a timeframe that is interpretable for epidemiological investigations of within- and between-farm transmission. Briefly, for each variant in the time-scaled trees, we extracted the time to the most recent common ancestor, which was used to calculate clade age, divergence time from the most closely related variant, and clade purity.

### Working group and practitioner survey

A working group was established in March, 2021 that included representatives from major swine-oriented diagnostic labs (University of Minnesota, Iowa State University, South Dakota State University, Ohio Animal Disease Diagnostic Lab), swine disease monitoring programs that serve as national repositories of PRRSV sequences (Morrison Swine Health Monitoring Program (27), Swine Disease Reporting System (35, 36), and USDA-Agricultural Research Service’s Swine Pathogen Database (37)), and swine veterinarians and production systems. This group was involved iteratively in the development of the new classification system (Supplementary Figure S2).

The working group developed and administered a survey to swine health professionals in the U.S. to better understand how they use genetic sequence data. In this survey, practitioners were asked to rank the top four reasons for which they submitted samples for sequencing (out of 10 options related to within-farm monitoring, between-farm or regional spread, or immunological/phenotype considerations). This survey was distributed in April 2022 by the American Association of Swine Veterinarians. Based on the results of this survey and with input from the working group, the final classification system was tailored to meet the needs of practitioners by explicitly addressing the primary motivations for sequencing.

## Acknowledgments

This study was funded by the joint NIFA-NSF-NIH Ecology and Evolution of Infectious Disease award 2019–67015-29918, the Intramural Research Program of the U.S. Department of Agriculture, National Institute of Food and Agriculture, Data Science for Food and Agricultural Systems Program, grant number 2023-67021-40018, a grant from the American Association of Swine Veterinarians, and the U.S. Department of Agriculture, Agricultural Research Service project 5030-32000-231-000-D. MSHMP was funded by the Swine health Information Center (SHIC, www.swinehealth.org, Project # 23-079).

The authors would like to thank members of the AASV PRRSV nomenclature working group, including Andreia Arruda, Srijita Chandra, Eric Nelson, Tom Petznick, Melanie Prarat, Mark Schwartz, Donna Drebes, Mark Wagner, Jessica Seate, Gustavo Silva, Joel Sparks, and Paul Yeske. The authors would like to thank the Morrison Swine Health Monitoring Project (MSHMP) participants and the Veterinary Diagnostic Laboratory, University of Minnesota, for sharing its PRRSV-2 genetic sequences. Mention of trade names or commercial products in this article is solely for the purpose of providing specific information and does not imply recommendation or endorsement by the U.S. Department of Agriculture (USDA). The funders had no role in study design, data collection and interpretation, or the decision to submit the work for publication. The findings and conclusions in this publication are those of the authors and should not be construed to represent any official USDA or U.S. Government determination or policy. USDA is an equal opportunity provider and employer.

Author order of the first three and last two authors were based on effort. The remaining authors were listed in reverse alphabetical order. KV designed the research, performed the analysis, and wrote the paper. NP, MK, and IP performed some analyses, contributed to interpretation, and edited the paper. NP and DN developed analytical pipelines and the RShiny app. DL, GT, JZ, TA, MZ, CC and DH contributed to research design, results interpretation, and manuscript editing.

