## Supplementary for "PRRSV-2 variant classification: a dynamic nomenclature for enhanced monitoring and surveillance"

### Supplementary material

**Supplementary Figure 1.** a) Mean within-variant genetic distance since year of emergence, and b) clade purity over time.

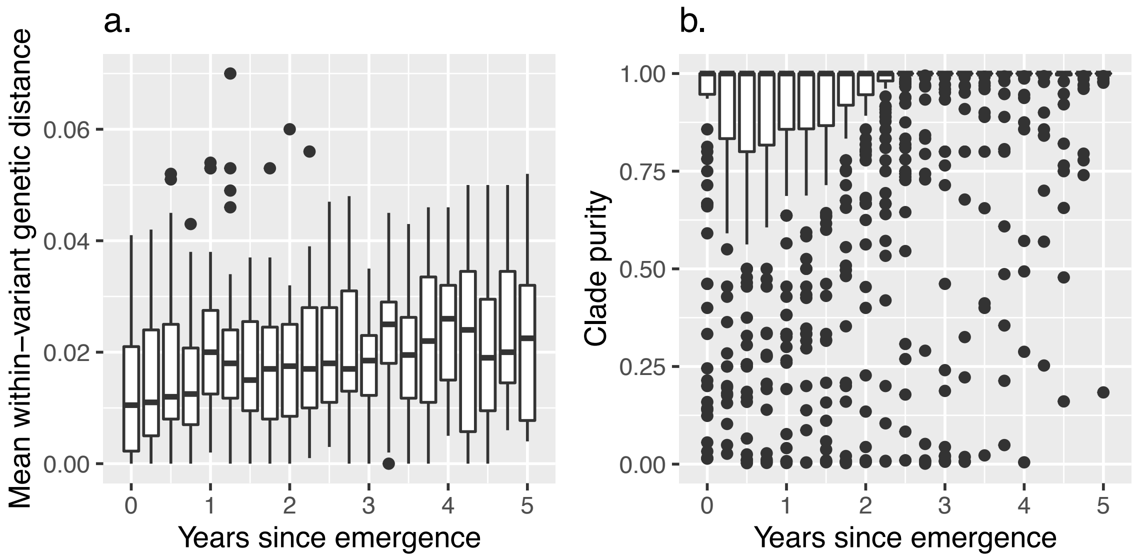


**Supplementary figure 2.** Schematic of iterative input solicited from working group for the development and refinement of the variant classification system.

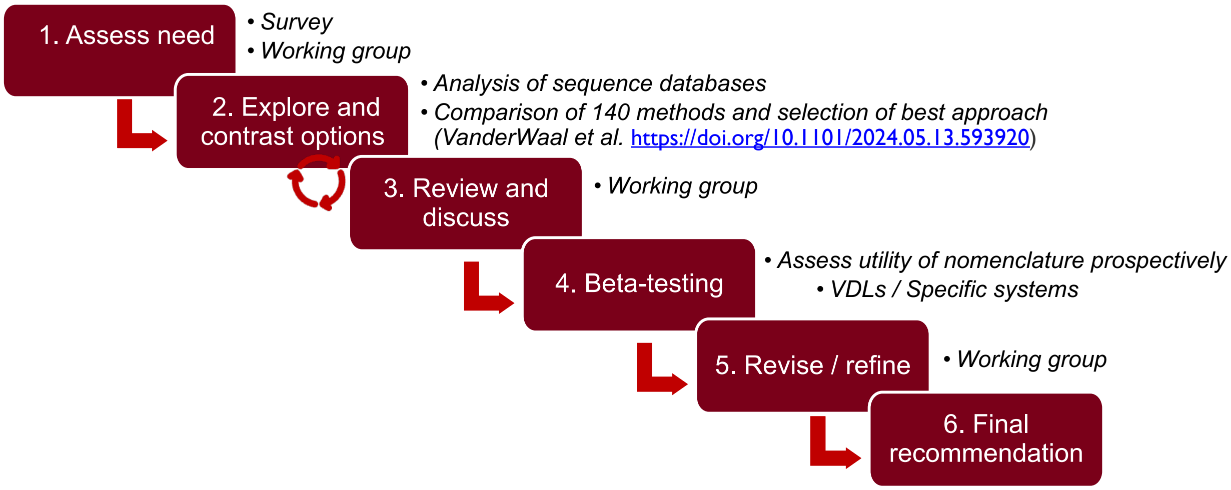


### Supplementary text

#### Post-processing of *TreeCluster* outputs

Additional steps were applied to *TreeCluster* outputs based on preliminary results showing that some variants defined on the first 36-month tree did not consistently group together in subsequent trees. Two reasons for the inconsistency of clustering stems from the way that *TreeCluster* handles a) inherent uncertainties in tree topology, and b) closely related clusters. *TreeCluster* attempts to handle topological uncertainties through a support argument, whereby all internal nodes within the cluster must have a bootstrap that is greater than a user-set threshold. However, sequences within a cluster have high nucleotide identity resulting in low bootstrap values for internal nodes. For our purposes, it was more important for the ancestral node of the clade to have high bootstrap support, thus supporting the existence of the clade overall regardless of its exact internal topology. *TreeCluster* does not evaluate support at the ancestral node. Thus, we post-processed *TreeCluster* outputs by extracting the ancestral bootstrap value for each cluster, as well as the identity and minimum genetic distance to the nearest neighboring cluster in the tree. Briefly, questionable clusters (with ancestral bootstrap values of <85 or distance to nearest neighbors of <0.02) were merged with their nearest neighboring clustering. This was repeated iteratively until there were no additional merges of questionable clusters possible. Some questionable clades could not be merged into well-supported clusters, and sequences belonging to such clades were listed as “unclassified.”

**Identifying new variants and splitting existing variants**

*Checking for new variants:* Phylogenetic clades were identified in the new tree using *TreeCluster* with post-processing, as described above, and the resulting clusters in the new tree were assigned temporary clade IDs. Because phylogenetic reconstructions are imperfect, not all sequences belonging to the same variant formed completely monophyletic groups when the trees were reconstructed on a quarterly basis. Thus, for each variant, the *TreeCluster* clades in the new tree were ranked according to what proportion of those clades belonged to that variant. The *TreeCluster* clades in a given tree were matched to the most likely variant ID following a greedy heuristic wherein the variant IDs were matched to the most likely clade (i.e., the clade in the tree that had the highest proportion of that variant ID), with more frequent variants being matched to the most likely clade first. The clades that best matched with each variant were identified by finding the clades that contained the highest proportion of a particular variant’s sequences. If two variants mapped to the same clade in the tree, then the clade was matched to the variant with the higher frequency, and the variant with the lower frequency was matched to its second most likely clade in the tree. Sequences that did not fall in the most likely clade of their assigned variant were flagged to be excluded from training dataset, and clades that were not matched to an existing variant ID were flagged as candidate new variants.

*Checking for splits in existing variants:* With the sequences from the most recent quarter added to the rolling database, the average pairwise genetic distance was calculated for each active variant based on the last 12 months of data. Variants where the 95$^{th}$ percentile of pairwise genetic distances was >5% were flagged as candidates to split. Sub-trees were created with the candidate variant, and divergent clades with ancestral branches with a length of >2%, bootstrap values of >85, and at least 5 descendent tips were identified as potentially needing to be split into a new variant. A new variant was only created if the minimum and median genetic distance between the new and original variant was >3% and >5%, respectively. This high threshold was set in order to minimize the number of sequences being re-named as a result of variant splitting, as per the request of diagnostic laboratories. Checking for splits based on the >5% genetic distance threshold was only performed for the quarters after December 31, 2023.

***Python assignment algorithm training***

With the same training dataset used in R, we fitted a random forest classifier for each quarter using the scikit-learn library in Python (Scikit-learn: Machine Learning in Python, (38)). Random search with ten-fold cross-validation was performed during the initial model training to tune the hyperparameters, ensuring optimal performance for subsequent quarters. These hyperparameters yielded the best classifier score of 0.97 while limiting the number of trees in the random forest (n_estimators) to 100, minimizing the model file size. The final random forest classifier model for the recent quarter is automatically downloaded from GitHub (https://github.com/kvanderwaal/prrsv2_classification) and performs variant assignment when the provided Python script is run locally. The assignment output format and nomenclature are consistent with the output from R's algorithms.

***Clade purity calculation***

Clade purity was calculated for each variant *j* in a sub-tree by first identifying the clade containing those sequences by finding the most recent common ancestor ($MRCA_j$) of all sequences belonging to that variant. We then identified all sequences descending from that ancestral node. Ideally, the descendent clade should purely contain sequences belonging to that variant; if sequences belonging to other variants were present within the descendent clade, this would indicate instability in the variant on the tree. We quantified the extent to which sequences belonging to other variants were present in variant *j*'s clade by calculating clade purity (proportion of sequences descending from the MRCA that belong to variant *j,* with 1 indicating perfect purity).

Clade purity was highly sensitive to single outlier sequences; if a single sequence is placed far away from the rest of the variant, then this results in a deep node being identified as the common ancestor, which means that a very large number of non-variant sequences are included in the clade, resulting in low purity metrics that are driven by a single outlier. To overcome the influence of outlier sequences, patristic pairwise distances were calculated between all members of a clade, and those with were in the 15% of sequences with the highest pairwise distance were excluded in the identification of the MRCA to avoid having a deep node being identified as the ancestor.

***Time-scaled trees***

We constructed time-scaled phylogenetic trees to contextualize the timeline of variant emergence and divergence on a timeframe that is interpretable for epidemiological investigations of within- and between-farm transmission. Given computational constraints, time-scaled trees were constructed separately for each sub-lineage using BEAST v.1.10.4 (75 sequences for L1BG, 568 for L1C, 795 sequences for L1H, and 129 for L1E/Dalpha sequences – note that sub-lineages were combined following Yim-im *et al.* 2023). The temporal signal and approximate time to the most recent common ancestor (tMRCA) of each sub-lineage was first estimated from maximum-likelihood trees using Tempest. The models were run with the GTR with invariant sites nucleotide substitution model, a lognormal uncorrelated relaxed molecular clock, and the Bayesian Skygrid population model with 50 parameters and the time at last transition point set to the estimated tMRCA from Tempest (or 10 years if the estimated tMRCA was <10 years). Duplicate MCMC chains of 100 million steps each, sampling every 10,000 steps. For L1A (the largest sub-lineage with 1675 sequences), sequences were down-sampled to 777 sequences using an in-house script that maintains genetic and temporal diversity (i.e., clades of closely related sequences were down-sampled to include only a single sequence per calendar quarter). In addition, only 25 Skygrid parameters were used for the L1A analysis to reduce computational burden and a chain length of 300 million was required for convergence. The first 10% of MCMC chains were discarded as burnin. The duplicate runs were inspected for convergence, and then combined using LogCombiner. Maximum clade credibility trees (MCC) were built using TreeAnnotator v.1.10.4 and visualized with *ggtree*.

The time to the most recent common ancestor (tMRCA) for each variant *i* was identified, and "clade age" was calculated as the average time elapsed between the $tMRCA_i$ and all sampled sequences belonging to that variant. The closest related variant was identified as the most recent ancestor of variant *i* whose descendants included sequences not belonging to variant *i*. Divergence time was calculated as the average time elapsed between this ancestor and all descendent sequences (inclusive of both variant *i* and related variants). Given that the short-term dataset included only three years of sequences, the analysis was repeated including all available sequences from the long-term dataset for one sub-lineage (L1H, 1702 sequences). Due to computational constraints, these data were down-sampled, maintaining genetic and temporal diversity. Clade age and divergence times were re-calculated from this tree and compared to the estimates from the short-term tree to assess the sensitivity of estimates to temporal scope.

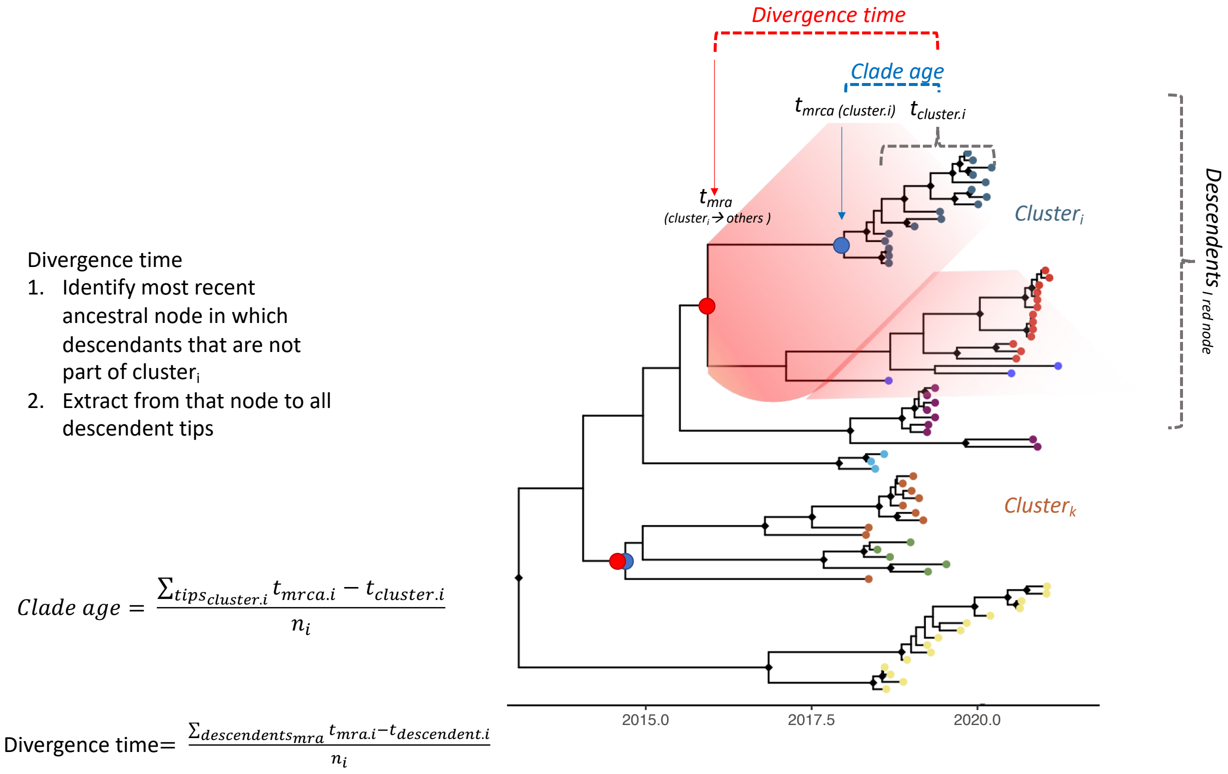
